# SVScore: An Impact Prediction Tool For Structural Variation

**DOI:** 10.1101/073833

**Authors:** Liron Ganel, Haley J. Abel, Ira M. Hall

## Abstract

**Motivation:** Structural variation (SV) is an important and diverse source of human genome variation. Over the past several years, much progress has been made in the area of SV detection, but predicting the functional impact of SVs discovered in whole genome sequencing (WGS) studies remains extremely challenging. Accurate SV impact prediction is especially important for WGS-based rare variant association studies and studies of rare disease.

**Results:** Here we present SVScore, a computational tool for *in silico* SV impact prediction. SVScore aggregates existing per-base single nucleotide polymorphism pathogenicity scores across relevant genomic intervals for each SV in a manner that considers variant type, gene features, and uncertainty in breakpoint location. We show that in a Finnish cohort, the allele frequency spectrum of SVs with high impact scores is strongly skewed toward lower frequencies, suggesting that these variants are under purifying selection. We further show that SVScore identifies deleterious variants more effectively than naïve alternative methods. Finally, our results indicate that high-scoring tandem duplications may be under surprisingly strong selection relative to high-scoring deletions, suggesting that duplications may be more deleterious than previously thought. In conclusion, SVScore provides pathogenicity prediction for SVs that is both informative and meaningful for understanding their functional role in disease.

**Availability:** SVScore is implemented in Perl and available freely at {{http://www.github.com/lganel/SVScore}} for use under the MIT license.

**Supplementary information:** Supplementary data are available at *Bioinformatics* online.

## 1 Introduction

Structural variation is an important source of human genome variation that includes deletions, duplications, inversions, mobile element insertions, translocations, and complex rearrangements. Over the past several years, much progress has been made in the area of structural variant (SV) detection, and we are now able to routinely detect 5,000-10,000 SVs in a typical deeply sequenced human genome (Sudmant *et al*., 2015a). However, predicting the functional impact of SVs discovered in whole genome sequencing (WGS) studies remains extremely challenging. Accurate SV impact prediction is especially important for WGS-based rare variant association studies and WGS-based studies of rare disease.

In recent years, there have been many efforts to predict the effects of single nucleotide polymorphisms (SNPs) *in silico*, including SIFT (Ng and Henikoff, 2001), PROVEAN (Choi *et al*., 2012), PolyPhen (Adzhubei *et al*., 2012), and VEP (McLaren *et al*., 2010). More recent methods such as fitCons (Gulko *et al*., 2015), CADD (Kircher *et al*., 2014), and Eigen (Ionita-Laza *et al*., 2016) precompute scores across the genome that predict the pathogenicity of a hypothetical variant at each locus. However, constructing similar methods to predict SV pathogenicity is more difficult due to the diversity of variant size and type. Variant type is important because, for example, a deletion spanning an entire gene is likely to have much different functional consequences than an inversion with the same breakpoint coordinates. Furthermore, current short-read sequencing technologies make precise SV breakpoint detection difficult, resulting in uncertainty regarding their exact location. SV impact prediction methods must take these all of these factors into consideration in order to robustly prioritize candidate pathogenic variants.

There have been a few attempts at SV impact prediction in the past. ANNOVAR (Yang and Wang, 2015) annotates large deletions and duplications that have been previously reported as well as naming genes affected by dosage-altering variants, but it does not make pathogenicity predictions, nor does it handle balanced rearrangements. VEP performs superficial consequence prediction for SVs, but only for a limited range of variant types (insertions, deletions, and duplications). Neither method provides a quantitative pathogenicity score.

## 2 Methods

We present SVScore, a novel computational tool for *in silico* SV impact prediction. SVScore depends on an existing set of per-base pathogenicity scores; here we use the precomputed SNP scores from CADD v1.3, although any other scoring scheme could potentially be used. For each SV in an SV callset in Variant Call Format (VCF), SVScore aggregates these scores across a set of genomic intervals determined by the variant type, affected gene features, and uncertainty in the location of the break-points (Figure 1). To aggregate these scores into interval scores, SVScore first uses tabix (Li, 2011) to extract the scores in the interval from a text file. If multiple scores are given for a single locus (e.g. CADD provides scores for all 3 possible nucleotide substitutions at each position), SVScore uses the maximum score per position. It then applies an operation (e.g. max or sum; See Operations) to each interval to summarize the per-base scores into interval scores. One score is computed for each interval-operation pair.

**Fig. 1.**
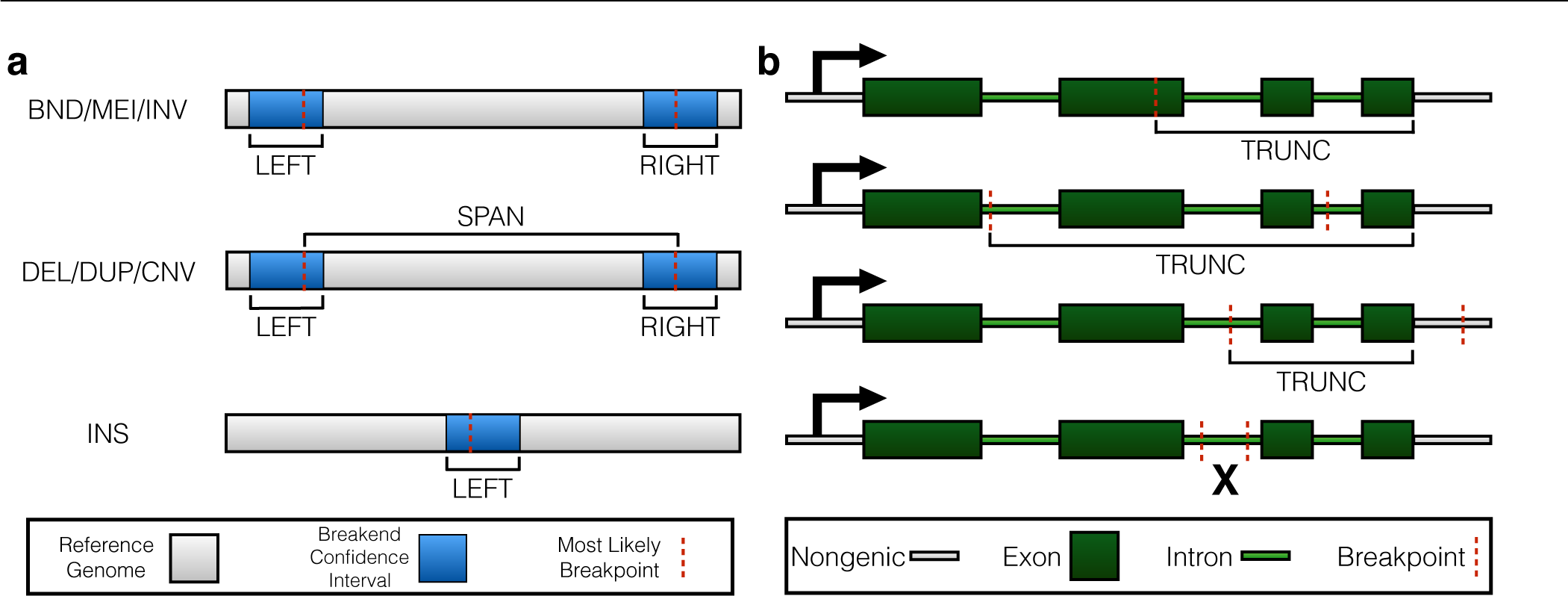
Intervals used by SVScore to aggregate per-base scores. **(a)** LEFT, RIGHT, and SPAN intervals chosen by SVScore based on SV type. LEFT and RIGHT scores comprise the entire confidence interval (CI) around the left and right breakpoint, respectively, and are calculated for every variant type. For deletions, tandem duplications, and other copy number variants (CNV), a SPAN score is calculated using the interval between the most likely breakpoints. LEFT and RIGHT scores are both calculated for novel sequence insertions (INS), but since these variants only have one breakpoint, the intervals and scores are the same for both sides. **(b)** Truncation scores (LTRUNC and RTRUNC) are calculated for those deletions, inversions, mobile element insertions, and novel sequence insertions that are predicted to truncate a transcript. The three cases in which these scores are calculated are: at least one break-point CI overlaps an exon, both breakpoint CIs are contained within different introns of the same transcript, and one breakpoint CI is contained within an intron while the other falls outside the transcript. In the case where both breakpoint CIs are contained within the same intron, no truncation score is calculated.

## 2.1 Intervals

As shown in Figure 1a, a score is calculated for dosage-altering variants over the interval between the most likely breakpoints (the boundaries of the SV-affected region), designated SPAN. The chosen operation is applied to the base scores in this region to calculate the SPAN score.

For every supported variant type, scores are calculated across the confidence intervals (CIs) around the left and right breakpoints. As insertions have only one breakpoint, the left and right CIs are the same. In these intervals (designated LEFT and RIGHT), the scores aggregated are the *possible breakpoint scores*, which are defined as the average of the scores of the 2 bases immediately flanking each possible breakpoint. If a breakpoint location is known precisely, the possible breakpoint score for this location is directly reported as the interval score regardless of the operation.

The final interval over which scores are calculated is the truncation interval, designated LTRUNC or RTRUNC depending on which break-point is involved. Truncation scores reflect the ability of certain SV types to truncate transcripts regardless of variant length (e.g. by disrupting an exon). The intervals across which these scores are calculated extend from the truncating breakpoint to the furthest downstream base of the affected transcript (Figure 1b). These scores are calculated for deletions, insertions, inversions, and mobile element insertions that intersect one or more genes. Figure 1b shows how SVScore decides which SVs are expected to truncate the transcripts that they overlap. Any breakpoint of a truncating type whose CI overlaps an exon is deemed to truncate the transcript. Furthermore, a variant whose breakpoint CIs are contained within two different introns is truncating, as is a variant with one CI in an intron and the other outside the transcript. Any variant whose break-point CIs are both completely contained within the same intron is not deemed truncating, so no truncation score is calculated for that transcript. SVScore uses vcfanno v0.0.11 (Pedersen *et al*., 2016) to find exons or introns that overlap SVs. As with SPAN scores, truncation scores are calculated from individual base scores rather than possible breakpoint scores.

## 2.2 Operations

The operations currently supported are: maximum, sum, mean, and mean of the top N scores. If multiple operations are selected in a single run, SVScore will apply all of the operations to each interval in parallel, reporting a score for each operation-interval pair. The maximum of all of a variant’s interval scores is reported as the score for the given operation and added to the INFO column of the VCF line(s). The individual interval scores can be reported as well using a command-line option.

SVScore supports weighting possible breakpoint scores using probability distributions calculated by tools such as LUMPY (Layer *et al*., 2014), as recorded in the INFO column. These give the probability of the true breakpoint being located at each possible breakpoint in a CI. Weighting the possible breakpoint scores using these distributions is important for two reasons. First, the expected score scales with size for the maximum and sum operations, causing a bias toward variants with large CIs. However, these variants are simply detected imprecisely, which is unrelated to their true pathogenicity. The second reason is that bases at a tail of the breakpoint probability distribution should not be given the same weight as those in the center of the distribution, as the former bases are less likely to be truly affected. When probability distributions are available, SVScore can incorporate them into the calculations of mean scores. If they are not present, SVScore simply assumes a uniform distribution over the CI. For weighted means of the top N bases in each interval, possible breakpoint scores are first weighted by the probability distribution, then the top N are chosen and the probability distribution over the chosen bases is rescaled to sum to 1.

Probability distribution weighting is only available when using the overall mean or the mean of the top N bases. Otherwise, weighting LEFT and RIGHT scores unfairly biases the scores toward dosage altering variants. These variants have a SPAN score that is unweighted by any probability distribution (as there is no probability distribution across a SPAN) and thus likely to be greater than LEFT and RIGHT scores of balanced rearrangements. However, the weighted mean of a breakpoint CI is similar in scale to the unweighted mean of a SPAN, making these comparisons fair.

## 3 Results and Discussion

To evaluate SVScore’s computational performance, we computed scores for a set of high confidence SVs called from WGS of ∼1,000 Finnish samples from an unrelated study (see Supplementary Methods). Scores were calculated using SVScore v0.5.1 with 5 operations – maximum, sum, weighted mean, and weighted mean of the top 10 and 100 bases in each interval. On a machine with two Intel Xeon E5-2670 processors (each with 16 threads) and 128 GB RAM, the total CPU time was 341 minutes. With 21,426 variants passing all of our filters, the average time per variant was 1.01 seconds. The average memory used was 1.7 GB, and the maximum memory used was 3.5 GB. Truncation scores, which are the most computationally expensive to calculate (due to intervals which may span large portions of transcripts) were calculated for 67.2% of variants in the SV callset (see Supplementary Figure 1 and Supplementary Table 1 for variant type composition and more detailed SVScore performance statistics).

In order to evaluate SVScore’s effectiveness in predicting deleterious variants, we used population allele frequency as a proxy for pathogenicity. Due to the effects of purifying selection, strongly pathogenic variants are likely to be observed at very low frequency in the human population. Thus, if SVScore is an accurate predictor of pathogenicity, the variants it predicts to be deleterious should be significantly more rare than those it predicts to be benign. For this experiment, impact scores were calculated using the weighted mean of the top 10 bases in each interval and exon/intron annotations from refGene. Figure 2 shows the allele frequency spectra of “pathogenic” variants (impact scores at or above the 90^th^ percentile), benign variants (impact scores below the 50^th^ percentile), and intermediate variants (all others). The pathogenic bin comprised 2 mobile element insertions, 561 tandem duplications, 1297 deletions, and 77 other novel adjacencies. (For this analysis, we excluded inversions, which comprised only 0.56% of the total variants detected.) These predicted pathogenic variants were heavily skewed toward the rare (AF < 0.01) end of the spectrum, while predicted benign variants were heavily skewed toward the common (AF >= 0.05) end, and variants with intermediate scores were between the other two categories. This suggests that high-scoring SVs are under strong purifying selection relative to low-scoring SVs, which strongly supports the utility of our impact scoring strategy.

**Fig. 2.**
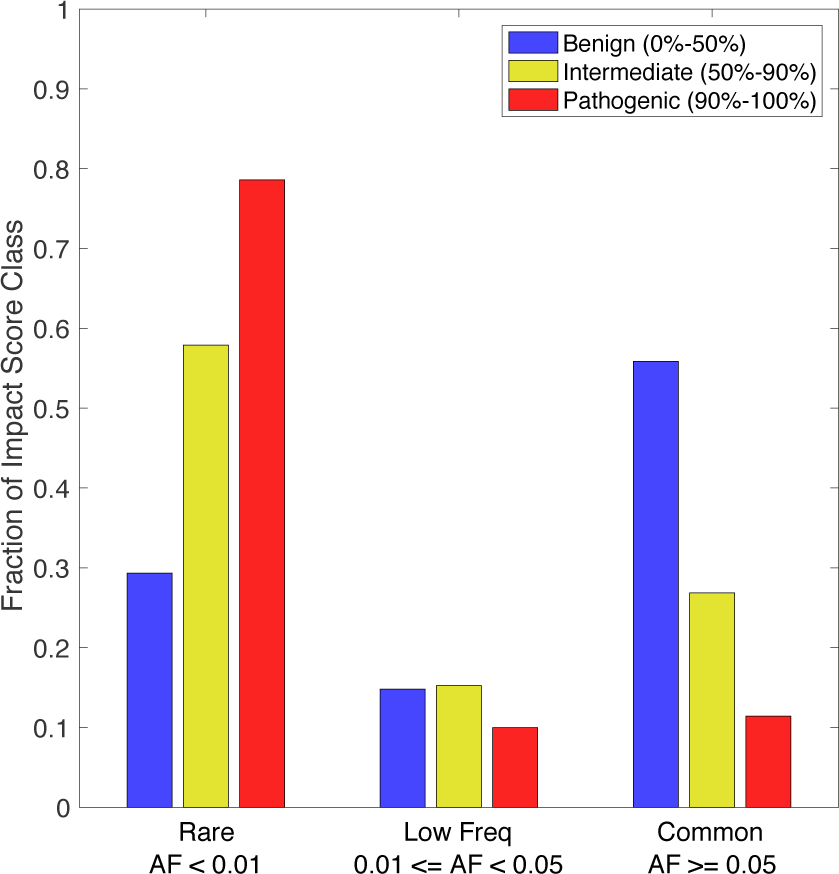
Allele frequency skew of high-scoring structural variants. Variants are separated first into classes based on impact score percentile, then into allele frequency bins. Each impact score class is normalized to 1 so that the height of each bar represents the fraction of the given impact score class that is in the given allele frequency bin.

To quantify the strength of this effect, we computed odds ratios representing how much more likely a predicted pathogenic variant is to be rare in a population than a predicted benign variant. Under the above definitions of benign, pathogenic, rare, and common, we found that pathogenic variants were significantly more likely than benign variants to be rare as opposed to common in the population (OR= 13.06, p = 5.43×10^-323^, Fisher’s Exact Test).

We calculated this odds ratio for several other definitions of “pathogenic” and “benign”, (Table 1, Figure 3). First, we tested several other SVScore percentile thresholds, keeping the definition of benign as the bottom 50% of variants (SVScore Threshold section of Figure 3). As the threshold is relaxed, the odds ratio generally decreases because increasing numbers of low-scoring variants are included in the pathogenic bin. The odds ratio of the top 1% of impact scores is very high – this is because at this threshold, there are 180 rare pathogenic SVs and only 12 rare benign ones. Next, we compared this to several odds ratios calculated for high confidence SNPs called on the same set of samples (see Supplementary Methods), using CADD v1.3 directly as the scoring scheme (SNP CADD Threshold section of Figure 3). To achieve similar odds ratios, we tested SNP thresholds that were much more stringent than the SV thresholds because the number of SNPs detected was several orders of magnitude greater than the number of SVs detected (with most SNPs expected to be benign). As with SVs, SNP odds ratios trend downward as the pathogenicity threshold is relaxed.

**Table 1.** Contingency table for SVs at varying impact score thresholds.

| SV Percentile | Rare | Common |
| --- | --- | --- |
| Bottom 50% | 2846 | 5420 |
| Top 1% | 180 | 12 |
| Top 5% | 772 | 119 |
| Top 10% | 1528 | 222 |
| Top 15% | 2212 | 351 |
| Top 20% | 2845 | 547 |
| Top 25% | 3449 | 754 |
| Top 30% | 3983 | 1032 |
| Top 35% | 4536 | 1294 |
| Top 40% | 5032 | 1623 |
| Top 45% | 5561 | 1934 |

We subsequently studied the variants in the top 10% of impact scores according to SVScore (Top 10% SVScores section of Figure 3, Supplementary Figure 2). To compare SVScore’s effectiveness in coding and noncoding regions, we calculated odds ratios for coding and noncoding SVs in this subset, again defining benign by the bottom 50% of all SV impact scores. All variants with at least one breakpoint CI or SPAN interval overlapping a refGene exon annotation were considered coding, and all others were designated noncoding. Coding SVs in the top 10% of impact scores had a greater odds ratio than noncoding variants in the same subset (13.68 for coding, 12.35 for noncoding), but the magnitude of this difference is surprisingly mild and suggests that many non-coding SVs are under similarly strong selection as coding SVs.

**Fig. 3.**
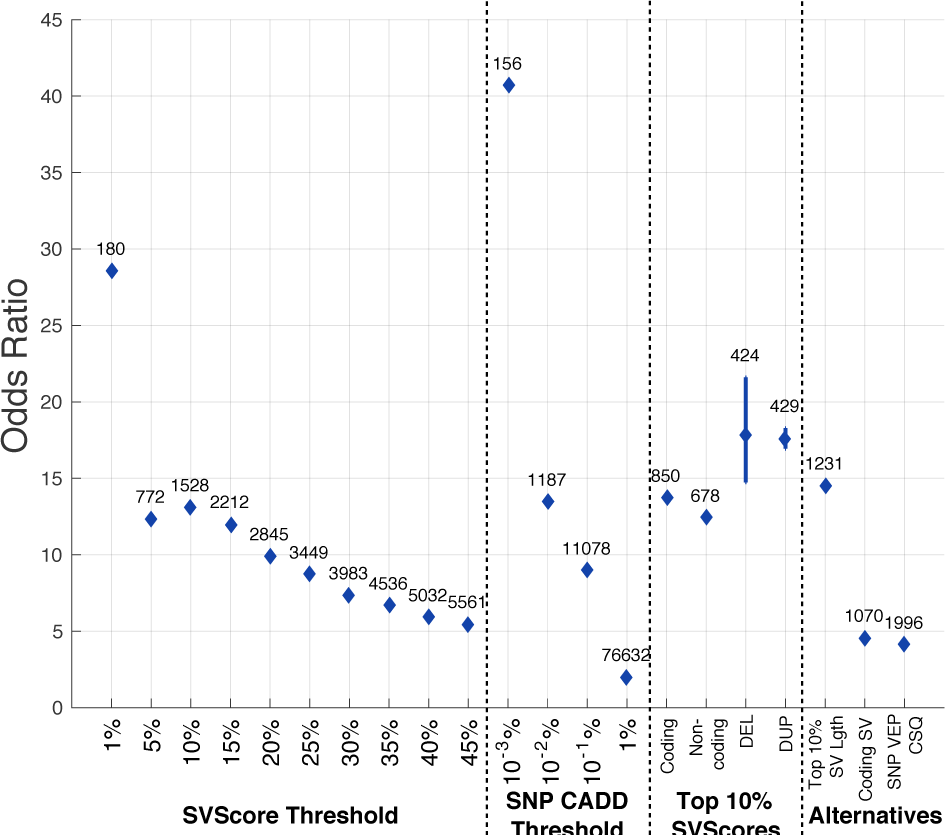
Allele Frequency Odds Ratio Comparison. Odds ratio is calculated as shown in equation (1). Above each point is the number of rare, pathogenic variants using the definition of pathogenicity given on the x-axis. The **SVScore Threshold** section shows the allele frequency odds ratios for SVs under varying definitions of pathogenicity based on impact score (top 1%, top 5%, etc.). For these odds ratios calculations, the variants in the bottom 50% of impact scores were considered benign. The **SNP CADD Threshold** section shows odds ratios calculated for SNPs using CADD at the thresholds shown (top 0.001%, top 0.01%, etc.). For these odds ratios, SNPs with CADD scores in the bottom 50% were used as benign variants. Pathogenic variants used for calculations in the Top 10% SVScores section were all subsets of those SVs with impact scores in the top 10%. In this section, the variants in the bottom 50% of all impact scores were again called benign. For the “Coding” and “Noncoding” experiments, the pathogenic variants were those SVs in the top 10% of impact scores that did and did not overlap a refGene exon, respectively. In the “DEL” and “DUP” experiments, the pathogenic variants were deletions and tandem duplications, respectively, in the top 10% of scores. The size distributions of these variants were matched as described in Supplementary Methods, and the 95% confidence intervals across the 100,000 samplings are shown. The **Alternatives** section shows three odds ratios from SVScore alternatives. In the “Top 10% SV Lgth” experiment, pathogenic variants were those with lengths at or above the 90^th^ percentile, and benign variants were those below the 50^th^ percentile of length. For “Coding SV”, pathogenic variants were those with at least one overlap between refGene exon and either breakpoint CI or SPAN interval, and benign variants were all others. Finally, the “SNP VEP CSQ” experiment used VEP’s IMPACT predictions for SNPs – variants with at least one HIGH prediction on a canonical transcript were called pathogenic, while those with only LOW or MODIFIER predictions on canonical transcripts were categorized as benign.

To compare the impact of high-scoring deletions and tandem duplications, we computed the same odds ratios for variants of these types within the top 10% of scores. Because duplications tend to be larger than deletions, we sampled SVs from both sets to make the size distributions approximately equal (see Supplementary Methods). Even when controlling for size in this way, the odds ratio for tandem duplications with impact scores in the top 10% was nearly equal to that for duplications (17.68 for deletions, 17.45 for duplications). This result may suggest that duplications are under much stronger selection than previously thought (Conrad *et al*., 2006; Cooper *et al*., 2011; Sudmant *et al*., 2015b). Alternatively, it is possible that this result reflects ascertainment bias against pathogenic deletions that cause embryonic lethality or severe developmental defects, and thus were not present in our adult Finnish cohort. Further work will be required to disentangle these factors.

To compare SVScore’s ability to distinguish high-impact variants to that of existing methods, we calculated odds ratios for two naïve alternatives (**Alternatives** section of Figure 3). First, we used SV length alone as a predictor of pathogenicity, categorizing large SVs (top 10% of lengths) as pathogenic and small SVs (bottom 50% of lengths) as benign. This yielded an odds ratio of 14.46, which is slightly greater than the odds ratio of 13.06 when using the top 10% of impact scores as “pathogenic”; however, substantially fewer rare, pathogenic variants were identified using the SV length method (1231 vs. 1538). While an SV’s length does influence its pathogenicity, calculated impact scores from SVScore are more effective predictors of pathogenicity than length alone. Figure 4 shows the size distributions for structural variants in our callset. As impact scores increase, the size distribution shifts toward larger variants. However, there is considerable overlap between the distributions, suggesting that while length is associated with pathogenicity, SVScore captures more information in its scores. Also, this method cannot be applied to translocations or other complex variants for which ‘length’ is not defined.

**Fig. 4.**
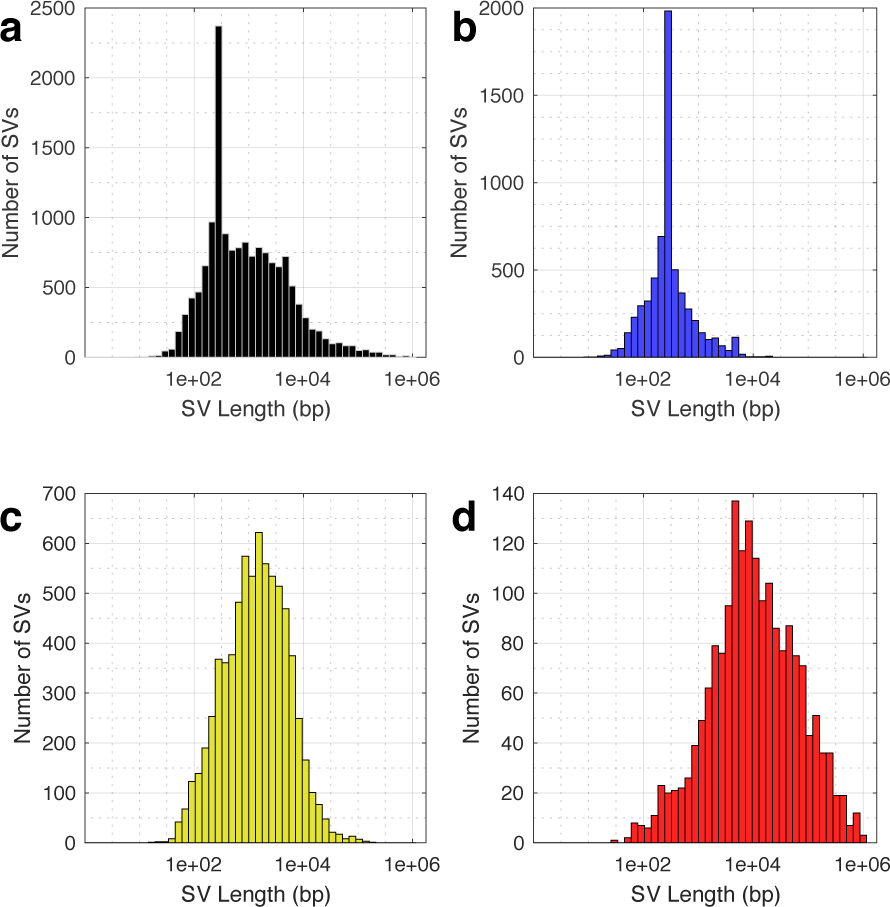
SV distributions separated by SVScore impact scores. SVs in each plot were placed into logarithmically sized length bins and plotted as histograms. **(a)** All SVs. **(b)** Benign SVs–impact scores below the 50^th^ percentile. **(c)** Intermediate SVs–impact scores below the 90^th^ percentile and at or above the 50^th^ percentile. **(d)** Pathogenic SVs – impact scores at or above the 90 percentile.

As a second naïve approach, we defined pathogenicity based on whether or not a structural variant (in any of its LEFT, RIGHT, or SPAN intervals) overlapped an annotated exon. This method identified fewer rare, pathogenic SVs (1070 vs. 1538) and resulted in a lower odds ratio than using the top 10% of impact scores.

We next sought to calibrate our SV impact scoring method with existing SNP-centric scoring methods. We first used IMPACT annotations from VEP to define pathogenicity of SNPs in our callset. SNPs with at least one IMPACT value of HIGH for a canonical transcript were categorized as pathogenic, while those with only LOW or MODIFIER values on canonical transcripts were designated benign. This approach was far less effective than SVScore in discriminating between pathogenic and benign variants, as evidenced by the lower odds ratio. Comparison of SVScore with CADD-based SNP impact scores revealed that the top 10% of highest scoring SVs (N=1,528) have a similarly strong allele frequency skew as the top 0.01% of SNPs (N=1,187). Interestingly, this result suggests that there may be a similar number of strongly pathogenic SVs and SNPs in the human population, despite the fact that SNPs are nearly 3 orders of magnitude more abundant overall.

## 4 Conclusion

SVScore is a novel *in silico* tool for predicting structural variant pathogenicity. In a large WGS dataset, predicted pathogenic variants were more depleted in a Finnish population than those of alternative methods. SVScore also identified pathogenic SVs in both coding and noncoding regions. While high-scoring variants tended to be longer, the length distributions of SVs in different score classes were not sufficiently different to justify using length alone to predict deleteriousness. Furthermore, we used SVScore to present evidence suggesting that tandem duplications are under similar levels of negative selection as deletions even when controlling for size, although further work is needed to confirm this.

We believe that SVScore will be a useful tool for future WGS-based studies by enabling facile prioritization of structural variants based on their likelihood of being deleterious. Its support for various operations and arbitrary per-base scoring schemes make it a powerful and flexible asset to investigators interested in the genomic alterations underlying both Mendelian and complex phenotypes.

## Acknowledgements

We thank Dave Larson and Indraniel Das for generating the SNP callset, Brent Pedersen for modifying vcfanno to support SVs, and members of the FinMetSeq consortium for contributing WGS data.

## Funding

This work has been supported by the NIH/NHGRI (5U54HG003079 and 1UM1HG008853) and a Burroughs Wellcome Fund Career Award (I.M.H).

## Conflict of Interest

none declared.

